# Inducible co-stimulatory molecule (ICOS) alleviates Paclitaxel induced peripheral neuropathy via an IL-10-mediated mechanism in female mice

**DOI:** 10.1101/2022.11.14.516419

**Authors:** Ishwarya Sankaranarayanan, Diana Tavares-Ferreira, Juliet Mwrigi, Galo Mejia, Michael D. Burton, Theodore J. Price

## Abstract

Chemotherapy-induced peripheral neuropathy (CIPN) is a primary dose-limiting side effect caused by antineoplastic agents, such as paclitaxel. This causes damage to peripheral nerves and the dorsal root ganglia (DRG). Currently, there are no effective treatments for CIPN, which can lead to long-term morbidity in cancer patients and survivors. Neuro-immune interactions occur in CIPN and have been implicated both in the development and progression of the disease and disease resolution. We investigated the potential role of Inducible co-stimulatory molecule (ICOS) in the resolution of CIPN pain-like behaviors in mice. ICOS is an immune checkpoint molecule that is expressed on the surface of activated T cells and promotes proliferation and differentiation of T cells. We found that intrathecal administration of ICOS agonist antibody (ICOSaa) alleviates mechanical hypersensitivity caused by paclitaxel and facilitates the resolution of mechanical sensitivity in female mice. Administration of ICOSaa reduced astrocyte-gliosis in the spinal cord and satellite cell gliosis in the DRG of mice previously treated with paclitaxel. Mechanistically, ICOSaa intrathecal treatment promoted pain resolution by increasing interleukin 10 (IL-10) expression in the dorsal root ganglion. In line with these observations, blocking IL-10 receptor (IL-10R) activity occluded the effects of ICOSaa treatment on CIPN behavior in female mice. Suggesting a broader activity in neuropathic pain, ICOSaa also partially resolved mechanical hypersensitivity in the spared nerve injury (SNI) model. Our findings support a model wherein ICOSaa administration induces IL-10 expression to facilitate neuropathic pain relief in female mice. ICOSaa treatment is in clinical development for solid tumors and given our observation of T cells in the human DRG, ICOSaa therapy could be developed for combination chemotherapy - CIPN clinical trials.

**Highlights:**

- ICOS agonist antibody (ICOSaa) promotes pain resolution in female mice
- DRG T cells appear to enter an anti-inflammatory phenotype by ICOSaa treatment
- ICOSaa treatment increases DRG levels of IL-10 cytokine
- ICOSaa effects in female mice are blocked by IL-10 sequestering treatment

## 1. Introduction

Chemotherapy-induced peripheral neuropathy (CIPN) is a debilitating condition due to the dose-limiting side effects of antineoplastic agents. Thirty-70% of patients receiving chemotherapy treatment experience numbness, reduced proprioception, and pain in their extremities [49]. CIPN is predominately sensory-related, causing damage to the peripheral nervous system with some motor deficits [6; 20]. Paclitaxel is a chemotherapy agent primarily used to treat breast, lung, and ovarian cancer. It acts through the stabilization of microtubules which leads to mitotic arrest [36].

Paclitaxel causes neurotoxicity by accumulating in the dorsal root ganglia (DRG) and in the peripheral nerves impairing axonal transport and causing mitochondrial dysfunction that is linked to the neuropathy [1; 12; 13; 15; 38; 39; 42; 47]. It is now well-accepted that the immune system can play an active role in modulating chronic pain progression and resolution [24; 55]. Neuro-immune interactions occur in CIPN, and both the adaptive and innate immune systems play an essential role in the progression and resolution of neuropathic pain [1; 19; 24]. For instance, immune cells have been shown to secrete both pro- and anti-inflammatory cytokines such as IL-6, IL-5, IL-4, IL-10, IFN γ and TNFα that can interact with neurons and regulate chronic pain [7; 30; 44; 63].

T cell activation occurs when the T cell receptor (TCR) interacts with major histocompatibility complex I and II expressed on antigen-presenting cells (APC), dendritic cells (DC), macrophages, and B cells; however, secondary signaling is required for proper facilitation of T cell activation [60]. The CD28 co-stimulatory family of receptors provides this secondary signal for the activation and survival of T cells. Inducible co-stimulatory molecule (ICOS) is a member of the CD28 family, and an immune checkpoint receptor expressed on activated T cells [2]. It is known to induce an immune response by binding to its exclusive ICOS ligand (ICOSL) expressed on APC, DC, B cells, and tumor cells [60]. ICOS, upon activation, generates a signaling cascade on various subsets of T cells such as the CD4 T helper cells and CD8 cytotoxic T cells [2; 3]. ICOS induction enhances proliferation and differentiation of T cells with the secretion of cytokines such as IL-4 and IL-10 [3; 33]. This makes ICOS-ICOSL an attractive therapeutic target for cancer immunotherapy [3] and neuropathic pain.

Previous studies have demonstrated that cytokine signaling is an important driver of CIPN, and this is a primary mechanism of neuro-immune modulation [7; 24; 39].

Paclitaxel promotes an increase in pro-inflammatory cytokines such as TNFα and IL1β with the suppression of anti-inflammatory cytokine IL-10 and IL-4 [28; 51; 57]. IL-10, an anti-inflammatory cytokine, exerts a neuroprotective and pain-relieving effect in CIPN, osteoarthritis, and chronic constriction injury-induced neuropathic pain [22; 40; 59]. IL-10 is secreted by innate immune and adaptive immune cells during various challenges [18; 41]. The IL-10 receptor (IL-10R) is expressed in DRG neurons where its activation suppresses the excitability of nociceptors providing a plausible cellular mechanism for the relief of pain promoted by IL-10 [21; 22; 45; 50; 61]. Based on this foundation of evidence, we hypothesized that activation of ICOS signaling on T cells in the DRG, could lead to the secretion of IL-10 cytokine and alleviate paclitaxel-induced peripheral neuropathic pain.

Our primary goal was to determine if ICOSaa treatment could promote pain resolution in paclitaxel-induced peripheral neuropathy in mice. We found that paclitaxel leads to T cell infiltration in the DRG, which supports the idea of targeting ICOS signaling for pain resolution. Administration of ICOSaa attenuated hind paw hypersensitivity in female mice previously treated with paclitaxel. ICOSaa also reduced astrogliosis in the dorsal horn of the spinal cord and satellite cell gliosis in the DRG. ICOSaa treatment led to enhanced expression of anti-inflammatory cytokine IL-10 in the DRG. Consistent with this, IL-10 receptor blocking treatment occluded the beneficial effect of ICOSaa treatment in the CIPN model. Our findings demonstrate a new mechanism for stimulating T cells to promote pain resolution via ICOSaa treatment.

## 2. Materials and Methods

### 2.1 Animals

ICR mice were maintained and bred at the animal facility at the University of Texas at Dallas. Experiments were performed using 8-12 weeks old female and male littermates. The mice were housed (4 maximum/cage) with food and water *ad libitum* in a 12h light-dark cycle and maintained at room temperature (21 ± 2°C). All procedures were approved by the Institutional Animal Care and Use Committee at University of Texas at Dallas.

### 2.2 Injections

Paclitaxel was dissolved in a mix of 50% El Kolliphor EL (Sigma-Aldrich) and 50% ethanol and diluted in sterile PBS. Mice received 4 mg/kg of paclitaxel every other day, for a cumulative intraperitoneal dosage of 16 mg/kg, or vehicle control (50% Kolliphor EL and 50% ethanol diluted in sterile PBS). Intrathecal injections of Inducible co-stimulatory (ICOS) agonist antibody (C398.4 Biolegend, 0.5 µg/µl) in total volume of 5 µl volume or PBS as vehicle control was administered using a 30-gauge (0.5’’) needle and Hamilton syringe for four consecutive days under isoflurane anesthesia starting from day 8 after the last dosage of paclitaxel. A tail flick was observed in mice as evidence of successful entry into the intradural space for intrathecal injections [16]. For the IL-10 receptor blocking treatment, mice received intraperitoneally 250 µg of InVivoMAb anti-mouse IL-10R antibody (BioXcell CD210) or InVivoMAb rat IgG1 isotype control, anti-horseradish peroxidase antibody (BioX cell) vehicle control twice weekly during the entire course of the experiment.

### 2.3 Mechanical withdrawal threshold

Mice were habituated for 1 hour before testing for mechanical hypersensitivity in a clear acrylic behavioral chamber. Mechanical paw withdrawal threshold was tested using the up-down method [8] using calibrated von Frey filaments (Stoelting) perpendicular to the mid plantar surface of the hind paw. A positive response consisted of an immediate flicking or licking behavior upon applying the filament to the hind paw. The investigator was blinded to treatment conditions during all days of testing.

### 2.4 Flow cytometry Analysis

Mice were euthanized under isoflurane anesthesia by cervical dislocation on day 13 after intrathecal injection of ICOSaa. L3-L5 DRGs were dissected and minced using scissors in the lysis buffer containing 1.6 mg/mL collagenase (Worthington), 10 mM HEPES (Thermo Fisher Scientific), and 5 mg/ml of Bovine serum albumin (Thermo Fisher Scientific) at 37ºC for 30 minutes followed by a gentle trituration. The digested tissue was passed through a 70 µm cell strainer, and the samples were centrifuged at 600×g at 4ºC for 5 min. The supernatant was discarded, and the cell pellet was incubated with red blood cell (RBC) lysis buffer (Biolegend) for 5 mins at room temperature followed by a wash with 1% Bovine serum albumin in Phosphate buffer saline (PBS) (Thermo Fisher Scientific). The resuspended cell pellet was stained for anti-mouse CD3-PE (1:200, 145-2C11 Biolegend), anti-mouse CD45-APC/Cy7 (1:200, 30-F11 Biolegend), anti-mouse CD8a-PE/Cy7 (53-6.7 Biolegend), anti-mouse CD4-PE-Cyanine 5.5 (RM4-5 Biolegend) in 1% BSA-PBS for 30 mins in the dark at 4ºC followed by three washes with 1% bovine serum albumin in PBS. The cell samples were acquired using BD LSR Fortessa (BD Biosciences) flow cytometer using BDFACS DIVA software (BD Biosciences) and analyzed with FlowJo software (v.10, FlowJo). Flow cytometry analysis was performed by first gating for CD45 positive hematopoietic cells followed by gating for single cell lymphocytes. We did not perform live/dead cell gating but all T cells were identified as CD45^pos^+CD3^pos^, subsets of T helper cells as CD45^pos^+CD3^pos^+CD4^pos^ and cytotoxic T cell subset as CD45^pos^+CD3^pos^+CD8a^pos^ [35].

### 2.5 Immunohistochemistry

Mice were euthanized under isoflurane anesthesia by cervical dislocation on day 13 after the last administration of ICOSaa. DRGs and spinal cord were dissected and fresh-frozen in optimum cutting temperature (OCT) medium (Fisher Scientific).

Human tissue issue procurement procedures were approved by the Institutional Review Boards at University of Texas at Dallas. Donor Information is provided in Table 1. Human dorsal root ganglion were collected within 4 hours of cross-clamp, frozen on dry ice and later embedded on OCT to cut on the cryostat. All tissues were sectioned at 20 µm using a cryostat (Leica) and placed on the SuperFrost charged side of the slides. The tissue sections were fixed in 4% cold formaldehyde (Thermo Fisher Scientific) for 10 minutes, followed by dehydration with increasing ethanol concentrations from 50%, 75%, and 100% for 5 minutes each. The sections were blocked for one hour using 10% normal goat serum (R&D systems) with 0.3% Triton-X100 (Sigma-Aldrich). The tissue sections were incubated with primary antibodies GFAP, Glutamine Synthetase, Peripherin, CD8a, CD4 (Table 2) diluted in blocking solution at 4ºC overnight, followed by the corresponding secondary antibody with DAPI diluted in blocking solution for 1 hour at room temperature. The slides were washed in 0.1 M PB and cover-slipped using Prolong Gold Antifade (Fisher Scientific P36930). Images were taken on Olympus FluoView 1200 confocal microscope or Olympus FluoView 3000 confocal microscope, using the same settings for all images per experiment.

**Table 1.** Human donor information

| Donor | DRG | Sex | Age | Cause of Death |
| --- | --- | --- | --- | --- |
| 1 | Lumbar | M | 65 | Stroke |
| 2 | Lumbar | M | 29 | Head Trauma |
| 3 | Lumbar | F | 57 | Stroke |
| 4 | Lumbar | F | 32 | Head Trauma |

**Table 2.** List of antibodies used.

| **Antibodies** | **Company** | **Catalog #** | **Dilution** |
| --- | --- | --- | --- |
| Glutamine Synthetase (GS) polyclonal antibody | Thermo fisher | 11307-2-AP | 1:1000 |
| Peripherin | EnCor Biotechnology Inc | CPCA- Peri | 1:1000 |
| DAPI | Cayman | 14285 | 1:5000 |
| Anti-GFAP | Neuro Mab | N206A/8 -75-240 | 1:1000 |
| Goat anti Mouse IgG (H+L) Secondary Antibody, Alexa Fluor 488, Invitrogen | Fisher Scientific | AA11029 | 1:2000 |
| IgG (H+L) Cross-Adsorbed Goat anti-Rabbit, Alexa Fluor® 555 | Fisher Scientific | A21428 | 1:2000 |
| IgY (H+L) Goat anti-Chicken, Alexa Fluor® 647 | Fisher Scientific | A21449 | 1:2000 |

Image analysis of the dorsal horn of the lumbar spinal cord were obtained by calculating the corrected total cell fluorescence (CTCF) intensity of GFAP using the formula CTCF = Integrated Density – (Area of selected cell × Mean fluorescence of background readings). CTCF values were normalized by the area the images captured and analyses of spinal cord images were done using ImageJ version 1.48 (National Institutes of Health, Bethesda, MD). A total of 3 sections per animal were analyzed.

L3-L5 DRG images were taken using the Olympus FluoView 3000 confocal microscope using the same setting for all images. A region of interest (ROI) was drawn around individual neurons to measure the mean grey intensity value (MGI). Peripherin was used as a guide to identify neurons, but all neurons were included in the calculation. Background fluorescence was also measured using negative control subjected to blocking and secondary antibody with no primary. An average intensity was calculated after subtracting the MGI values of negative control and normalized over the area of the individual ROI. Image analysis was performed using Olympus cellSens software.

### 2.6 Enzyme-linked immunosorbent Assay (ELISA)

DRGs were dissected on day 13 after the last administration of ICOSaa, and flash frozen on dry ice. The frozen tissue was homogenized using a sonicator in lysis buffer (50□mM Tris, pH 7.4, 150□mM NaCl, 1□mM EDTA, and 1% Triton X-100) containing protease and phosphatase inhibitors (Sigma-Aldrich) and briefly centrifuged to extract the protein from the supernatant. Pierce BCA Protein Assay (Thermo Fisher Scientific) was performed by following the manufacturer’s protocol to identify the amount of protein in each sample. Enzyme-linked immunosorbent assay to detect I□-10 (Thermo Fisher Scientific-88-7105-22) cytokine was performed according to the manufacturer’s instructions.

### 2.7 Surgery

Spared nerve injury (SNI) was performed as previously described [11], sparing the sural branch and ligating and cutting the peroneal and tibial branches at the left sciatic nerve trifurcation. Sham control was performed the same way but without cutting any nerve. Mice were allowed to recover for two weeks post-surgery before administration of the ICOSaa treatment and testing for mechanical von Frey thresholds.

### 2.8 Data analysis and Statistics

All analyses and data were generated using GraphPad Prism 8.4.1. Statistical analysis between groups were determined using one or two-way ANOVA, followed by Bonferroni, Sidak or Tukey *post hoc* tests. Differences between two groups were assessed using Student’s T-test. Statistical results can be found in the figure legends. Effect sizes were determined by subtracting behavior scores starting with ICOSaa administration (day 8) from baseline measures. Absolute values were summed from the beginning of ICOSaa treatment (starting from day 8) and plotted for each group. All data were represented as mean +/-SEM with p<0.05 considered significant. The sample size and sex are noted in the graphs and figure legends.

## 3. RESULTS

### 3.1 ICOSaa accelerates the resolution of paclitaxel-induced peripheral neuropathy in female mice

T cells activation can occur with the engagement of ICOS similar to CD28 [31; 34]. For that reason, we first assessed the T cell response after paclitaxel administration using flow cytometry. We measured the number of CD3 positive T cells by gating for singlets with CD45 positive lymphocytes, followed by subsequent gating for CD3 positive cells (Fig 1A, B). We observed a significant increase in the percentage of CD3 positive T cells in the DRG day 13 after the start of paclitaxel treatment compared to vehicle control (Fig 1C). Having confirmed an increase in the number of T cells in the DRG in mice treated with paclitaxel, we then tested whether ICOSaa would have any effect on paclitaxel-induced mechanical hypersensitivity. To do this, we treated both male and female mice with paclitaxel every other day for a total of 4 injections and cumulative dose of 16 mg/kg, which produced mechanical hypersensitivity in both sexes (Fig 2A). On the final day of paclitaxel treatment, we started once daily treatment of ICOSaa (0.5 µg/µL) or vehicle given intrathecally for 4 consecutive days. Qualitatively, animals of both sexes treated with ICOSaa showed a more rapid resolution of mechanical hypersensitivity, but a significant treatment effect was only seen in female mice by two-way ANOVA (Fig 2B). There was a significant effect on the overall effect size of ICOSaa treatment in female mice (Fig 2C). While there was a trend to faster resolution of mechanical hypersensitivity in male mice treated with ICOSaa, the effect was not significantly different from vehicle either at individual time points (Fig 2D) or in effect size (Fig 2E). Given the clear effect seen in female mice of ICOSaa, and the far greater proportion of women treated with paclitaxel for cancer, we focused on the female mice for the remainder of the study.

**Figure 1:**
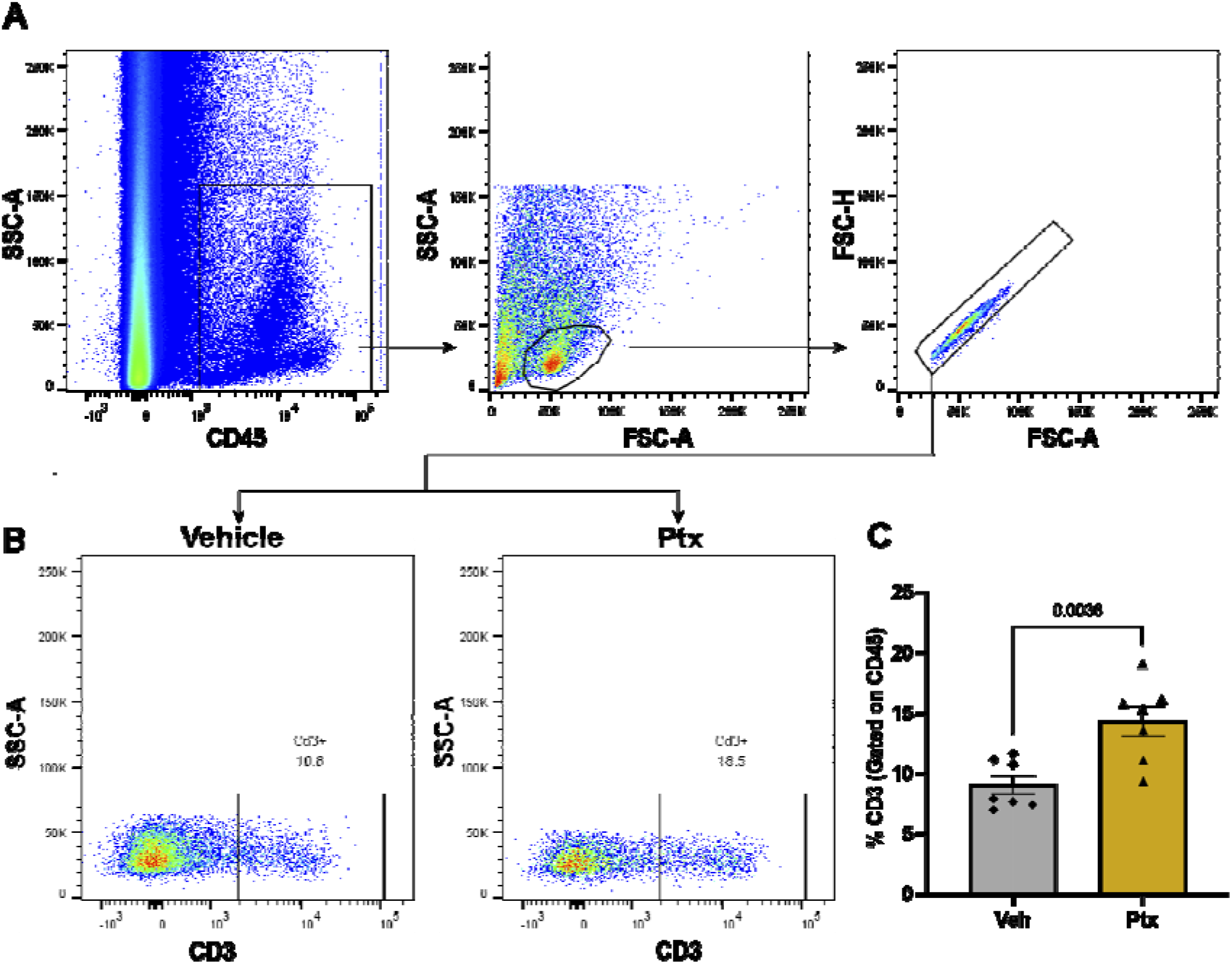
Paclitaxel treatment promotes infiltration of T cells into the DRG. A) Flow cytometry gating strategy for T cells in female mice gated for CD45^pos^ singlets isolated from L3-L5 DRG on day 13 after paclitaxel treatment. B) Representative flow cytometry plots for CD3^pos^ T cells (previously gated for CD45^pos^ singlets) on day 13 in mice treated with paclitaxel or vehicle C) Paclitaxel treatment was associated with a significant increase in the influx of T cells in the DRG measured by flow cytometry (unpaired t-test, t = 3.601, p-value = 0.0036, df=12) N=7/group.

**Figure 2:**
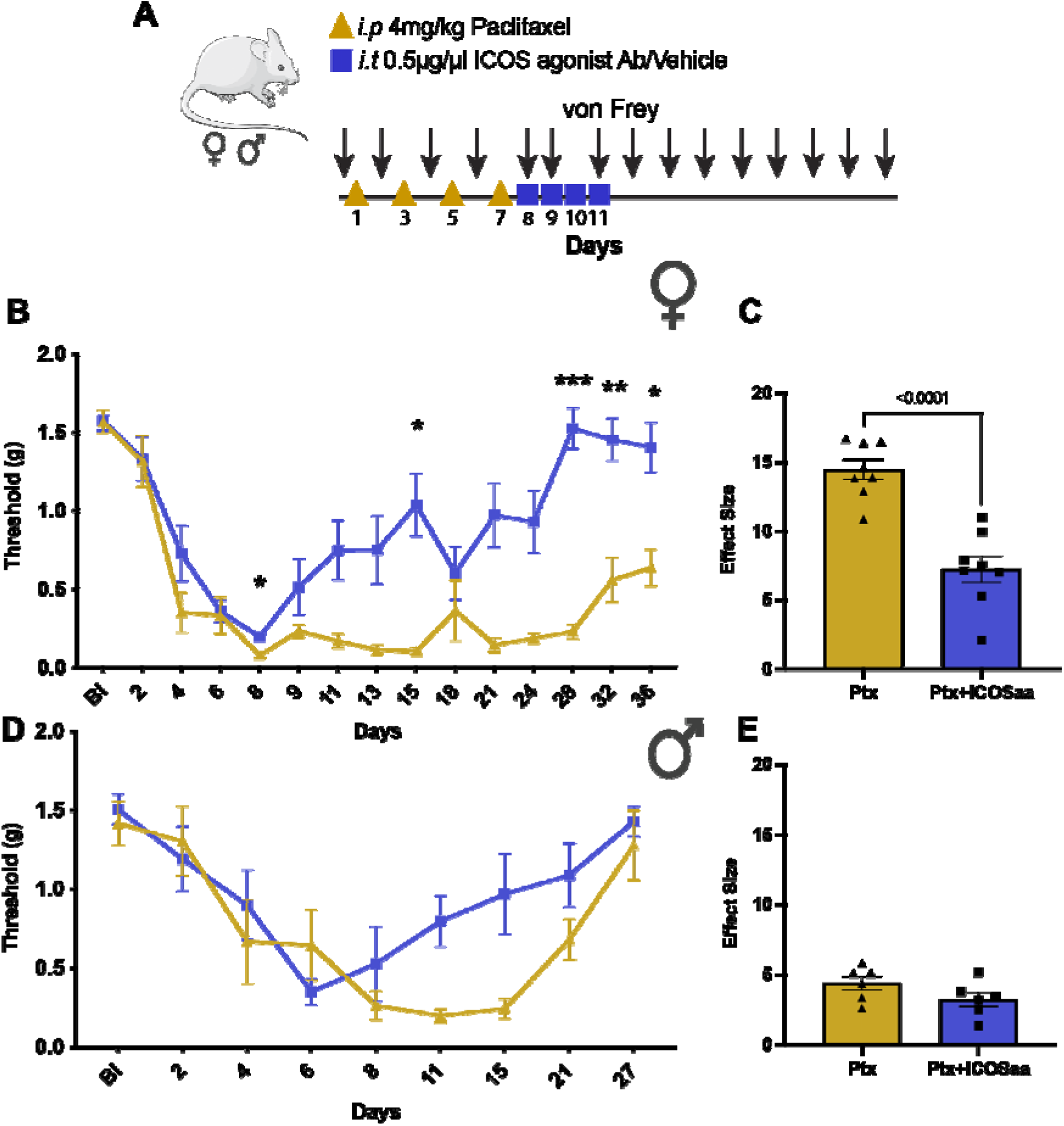
ICOS agonist antibody (ICOSaa) promotes the resolution of mechanical hypersensitivity in paclitaxel-induced peripheral neuropathy. A) The cohorts of mice were subjected to intraperitoneal injection of 4 mg/kg paclitaxel every other day for a cumulative dosage of 16 mg/kg according to the schema shown followed by intrathecal injection of ICOSaa or vehicle for four consecutive days. Arrows represent days of von Frey testing. Paclitaxel group represented in yellow and Ptx+ICOSaa represented in blue. B) Female mice reversed mechanical allodynia after intrathecal administration of ICOSaa (Two-way ANOVA, F=4.951, p-value<0.0001, post-hoc Sidak’s, Ptx+ICOSaa vs. Ptx, p-value=0.0318 at day 8, Ptx+ICOSaa vs. Ptx, p-value=0.0323 at day 15, Ptx+ICOSaa vs. Ptx, p-value=0.0001 at day 28, Ptx+ICOSaa vs. Ptx, p-value=0.0067 at day 32, Ptx+ICOSaa vs. Ptx, p-value=0.0.0261 at day 36), N=8/group C) Effect size was determined by calculating the cumulative difference between the value for each time point and the baseline value. The effect size difference was significant in the female cohort of mice (Effect size, unpaired t test, t = 4.963, p-value = 0.0002, df=14). D) Male mice showed a trend in resolution of mechanical hypersensitivity measured with von Frey filaments after administration of ICOSaa but it was not significant (Two-way ANOVA, F=2.020, p-value= 0.0544), N=6/group. E) We did not observe any statistically significant differences between the groups in male (Effect size, unpaired t test, t = 1.533, p-value = 0.1564, df=10).

### 3.2 ICOSaa reverses satellite cell gliosis in the DRG and astrogliosis in the spinal cord in paclitaxel treated female mice

Since we observed a resolution of mechanical allodynia in the ICOS-treated animals, we next examined glial cell changes in the DRG and the spinal cord [1; 29; 35; 46].

Gliosis is a sign of injury or stress to the tissue, where the cells become hyperactive and change morphology [43]. We performed immunohistochemistry to quantify the expression of glutamine synthetase [35] in satellite glial cells (SGC) in the DRG and GFAP in astrocytes in the dorsal horn of lumbar spinal cord. We observed an increase in Glutamine Synthetase (GS) expression in paclitaxel treated animals and a trend towards a reduction in animals treated with ICOSaa (Fig 3A, B). Expression of GFAP protein was increased in the dorsal horn of lumbar spinal cord in paclitaxel-treated animals and this was reduced in cohorts subjected to ICOSaa treatment (Fig 4A-C).

**Figure 3:**
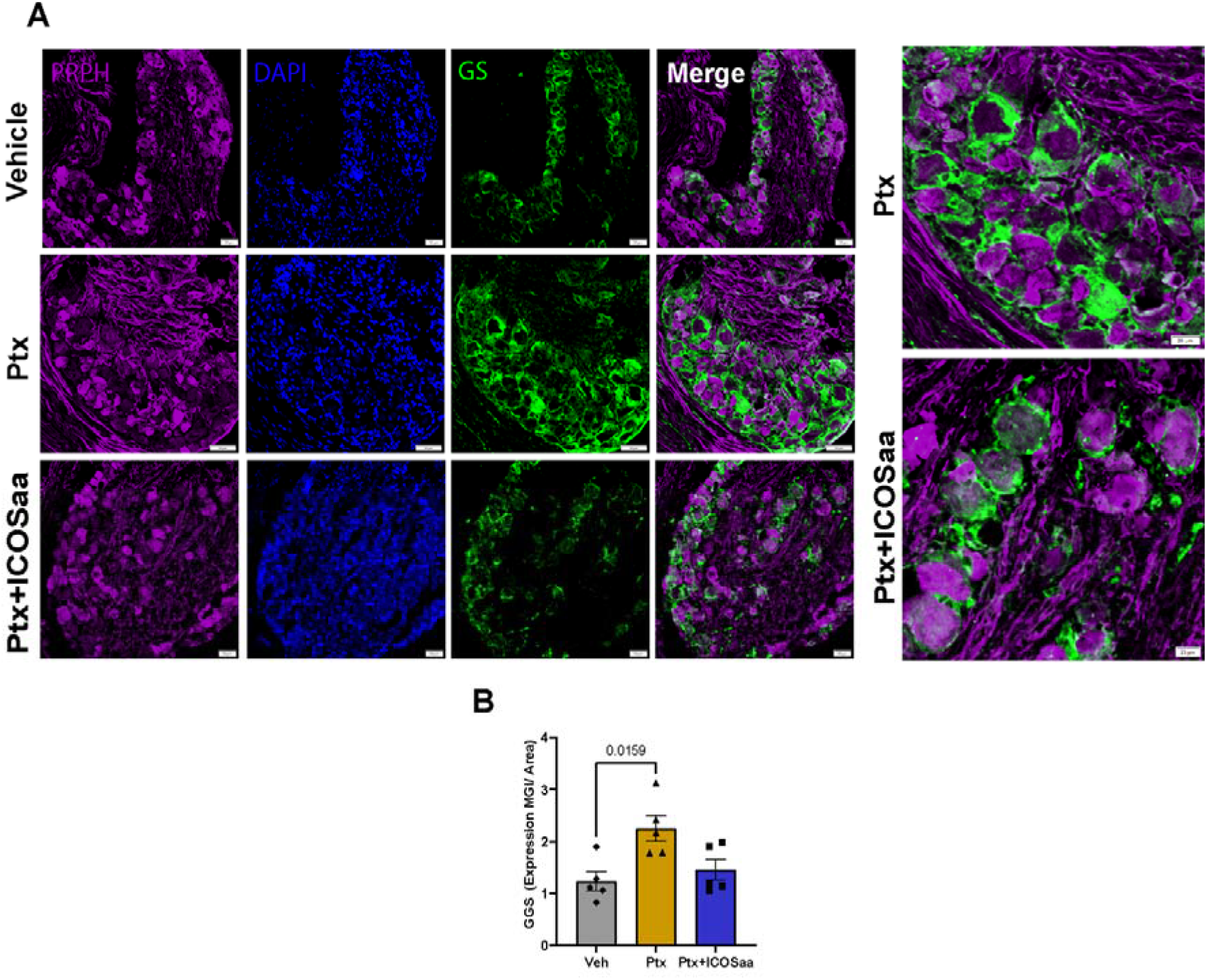
ICOSaa inhibits satellite cell gliosis in paclitaxel treated mice in the DRG. A) Representative images of GS that labels satellite glial cells (green) in the DRG, peripherin (purple) and Dapi (blue) B) GS trends towards a reduction in female animals treated with ICOSaa and calculated using MGI (One-way ANOVA, F=6.399, p-value=0.0128, post-hoc Tukey, Vehicle vs. Ptx, p-value=0.0159, Ptx vs. Ptx+ICOSaa, p-value= 0.0612) N=5/group. Data are represented as mean ± SEMs. Scale bar = 50µm.

**Figure 4:**
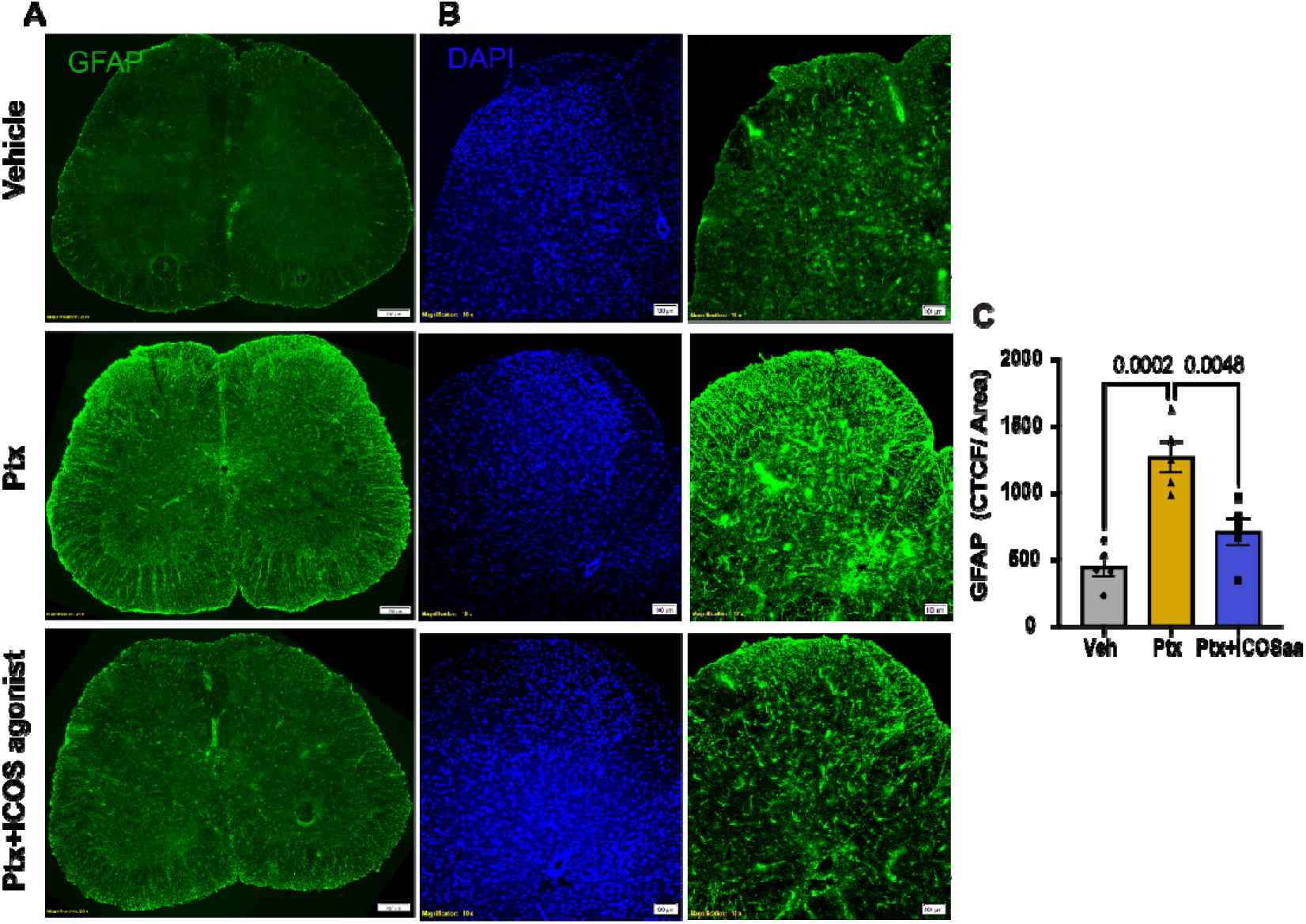
ICOSaa reverses astrocyte gliosis in the spinal cord in paclitaxel treated mice. A) Representative image of astrocyte labeling using GFAP antibody in the spinal cord of Vehicle, Ptx, and Ptx+ICOSaa groups from female mice. B) Representative images of the dorsal horn of the spinal cord of vehicle, Ptx, and Ptx+ICOSaa animal cohorts stained with GFAP (green) for astrocyte labeling and DAPI (blue). C) Paclitaxel animals showed increased expression of GFAP in the dorsal horn of the spinal cord. This was significantly reduced in mice treated with ICOSaa and calculated using CTCF (One-way ANOVA, F=18.42, p-value=0.0002, post-hoc Tukey, Vehicle vs. Ptx, p-value=0.0002, Ptx vs. Ptx+ICOSaa, p-value= 0.0048, Vehicle vs. Ptx+ICOSaa, p-value= 0.2485) N=5/group. Data are represented as mean ± SEMs. Scale bar = 100µm and 200µm respectively.

### 3.3 IL-10R antagonist blocks paclitaxel-induced mechanical hypersensitivity in mice treated with ICOSaa

Paclitaxel treated female mice that were then treated with ICOSaa showed a significant reduction in mechanical hypersensitivity. We hypothesized that this effect could be driven by IL-10 based on multiple studies demonstrating the T cell derived IL-10 can resolve neuropathic pain, in particular in female mice [21; 22; 26]. We injected IL-10R antagonist antibody or isotype control twice every week to block the activity of IL-10 at IL-10R, using the same paclitaxel and ICOSaa dosing schedule as described above (Fig 5A). We observed that while the IL-10R antagonist antibody had no effect on its own, it completely blocked the effect of ICOSaa treatment in paclitaxel treated female mice (Fig 5B, C). This finding suggests that the mechanism by which ICOSaa resolves paclitaxel-induced mechanical hypersensitivity is through the secretion of anti-inflammatory IL-10. Next, we assessed if IL-10 expression could be increased in the DRG after ICOSaa administration. We measured IL-10 concentration in the DRG on day 13, the day after the end of ICOSaa treatment, using an ELISA assay. We observed a significant increase in production of IL-10 cytokine in the ICOSaa treated cohort compared to paclitaxel or vehicle control (Fig 5D).

**Figure 5:**
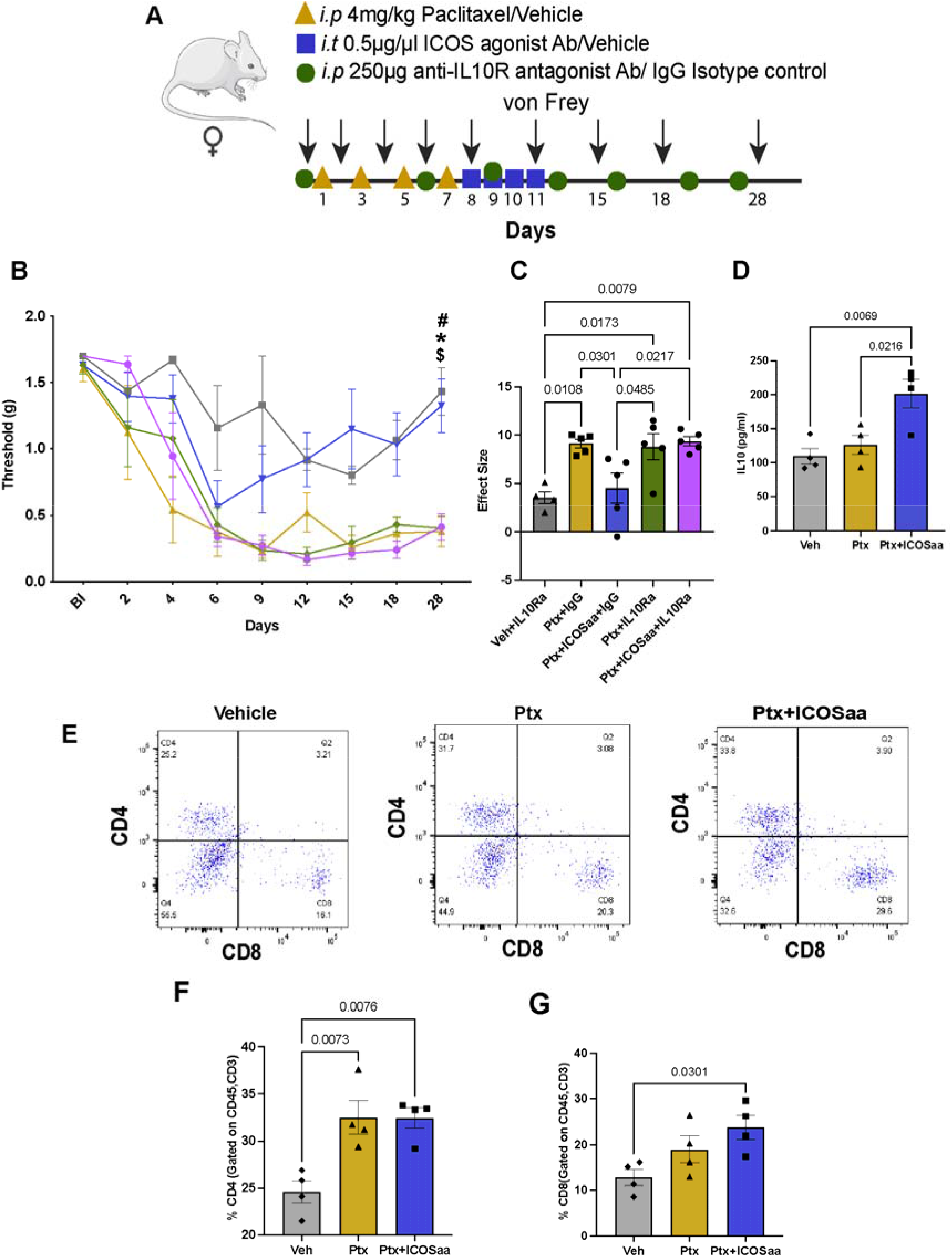
IL-10R antagonist reverses the effect of ICOSaa treatment in mice treated with paclitaxel. A) Schematic representation of experimental design. Arrow represents days with von Frey testing. Groups are shown as: Vehicle+ IL-10Ra (grey), Ptx+IgG (yellow), Ptx+ICOSaa+IgG (blue), Ptx+ICOSaa+IL-10Ra (pink), Ptx+IL-10Ra (green) B) Mechanical nociceptive thresholds are shown for each group with statistical differences represented in the graph (Two-way ANOVA, F=1.921, p-value=0.0048, post-hoc Bonferroni, Vehicle+IL-10Ra vs. Ptx+IgG, p-value=0.0362, Vehicle+IL-10Ra vs. Ptx+ICOSaa+IgG, p-value=ns, Vehicle+IL-10Ra vs. Ptx+IL-10Ra, p-value=0.0485, Vehicle+IL-10Ra vs. Ptx+ICOSaa+IL-10Ra, p-value= 0.0452, ^$^Ptx+IgG vs Ptx+ICOSaa+IgG, p-value= 0.0574, Ptx+IgG vs Ptx+IL-10Ra, p-value= ns, Ptx+IgG vs. Ptx+ICOSaa+IL-10Ra, p-value=ns, ^#^Ptx+ICOSaa+IgG vs. Ptx+IL-10Ra, p-value= 0.0697, *Ptx+ICOS+IgG vs. Ptx+ICOS+IL-10Ra, p-value= 0.0697, Ptx+ICOSaa+IL-10Ra vs. Ptx+IL-10Ra, p-value= ns). N = 4, 5, 5, 5, 5 per group, respectively. C) The effect size was significant between Ptx+ICOSaa+IL-10Ra and Ptx+ICOSaa+IgG1 (One-way ANOVA, F=7.211, p-value=0.0010, post-hoc Tukey, Vehicle+IL-10Ra vs. Ptx+IgG, p-value=0.0108, Vehicle+IL-10Ra vs. Ptx+ICOSaa+IgG, p-value=ns, Vehicle+IL-10Ra vs. Ptx+IL-10Ra p-value=0.0173, Vehicle+IL-10Ra vs. Ptx+ICOSaa+IL-10Ra, p-value=0.0079, Ptx+IgG vs. Ptx+ICOSaa+IgG, p-value=0.0301, Ptx+IgG vs. Ptx+IL-10Ra, p-value=ns, Ptx+ICOSaa+IgG vs.Ptx+IL-10Ra, p-value= 0.0485, Ptx+IL-10Ra+IgG vs.Ptx+ICOSaa+IL-10Ra, p-value= ns) D) A significant increase in IL-10 expression was observed in mice subjected to intrathecal injection of ICOSaa using ELISA (One-way ANOVA, F=9.452, p-value=0.0061, post-hoc Tukey, Vehicle vs. Ptx, p-value=ns, Vehicle vs. Ptx+ICOSaa, p-value=0.0069, Ptx vs. Ptx+ICOSaa, p-value= 0.0216) N=4/group E) Representative flow cytometry plots of subset of T cells CD4 positive and CD8 positive (previously gated on CD45^pos^CD3^pos^) in each group after treatment, isolated on day 13 from L3-L5 DRG. F) Percentage of CD4 positive T cells was increased in Ptx and Ptx+ICOSaa groups compared to vehicle treated cohort and no significant changes were observed between Ptx and Ptx+ICOSaa groups (One-way ANOVA, F=10.86, p-value=0.004, post-hoc Tukey, Vehicle vs. Ptx,p-value=0.0073, Vehicle vs. Ptx+ICOSaa, p-value=0.076, Ptx vs. Ptx+ICOSaa, p-value= ns) N=4/group G) Percentage of CD8 positive T cells were increased only in Ptx+ICOSaa group compared to vehicle control (One-way ANOVA, F=4.881, p-value=0.0367, post-hoc Tukey, Vehicle vs. Ptx, p-value=ns, Vehicle vs. Ptx+ICOSaa, p-value=0.0301, Ptx vs. Ptx+ICOSaa p-value= ns), N=4/group, *p < 0.05

Building on our observation that paclitaxel administration recruits T cells into the DRG, we sought to further examine the subsets of T cells infiltrating the DRG after administration of ICOSaa. We performed flow cytometry analysis by gating for T cell subsets as CD45^pos^CD3^pos^CD4^pos^ and CD45^pos^CD3^pos^CD8a^pos^ (Fig 5E). We saw no significant differences in the percentage of CD4 and CD8 positive T cells between ICOSaa treated cohort and paclitaxel treated mice (Fig 5F and G). We observed an increase in the frequency of CD8 positive T cells in ICOSaa mice. This cell type is known to promote upregulation of IL-10 in the DRG [21; 23]. This further suggests that ICOSaa either directly activates T cells or does so indirectly to increase endogenous IL-10 expression promoting the resolution of paclitaxel induced neuropathy in female mice.

### 3.4 ICOS agonist alleviates mechanical hypersensitivity in the SNI model

We next investigated if ICOSaa treatment could have an effect on mechanical hypersensitivity in the SNI neuropathic pain model. We performed SNI or sham surgery and administered four consecutive doses of ICOSaa two weeks post-surgery (Fig 6A). We observed an inhibition of mechanical hypersensitivity with ICOSaa treatment in female SNI mice on days 17 and 19 post SNI. This effect was transient and lasted only until the last dose of ICOSaa administration (Fig 6B). The effect size was also significant in animals treated ICOSaa compared to vehicle (Fig 6C).

**Figure 6:**
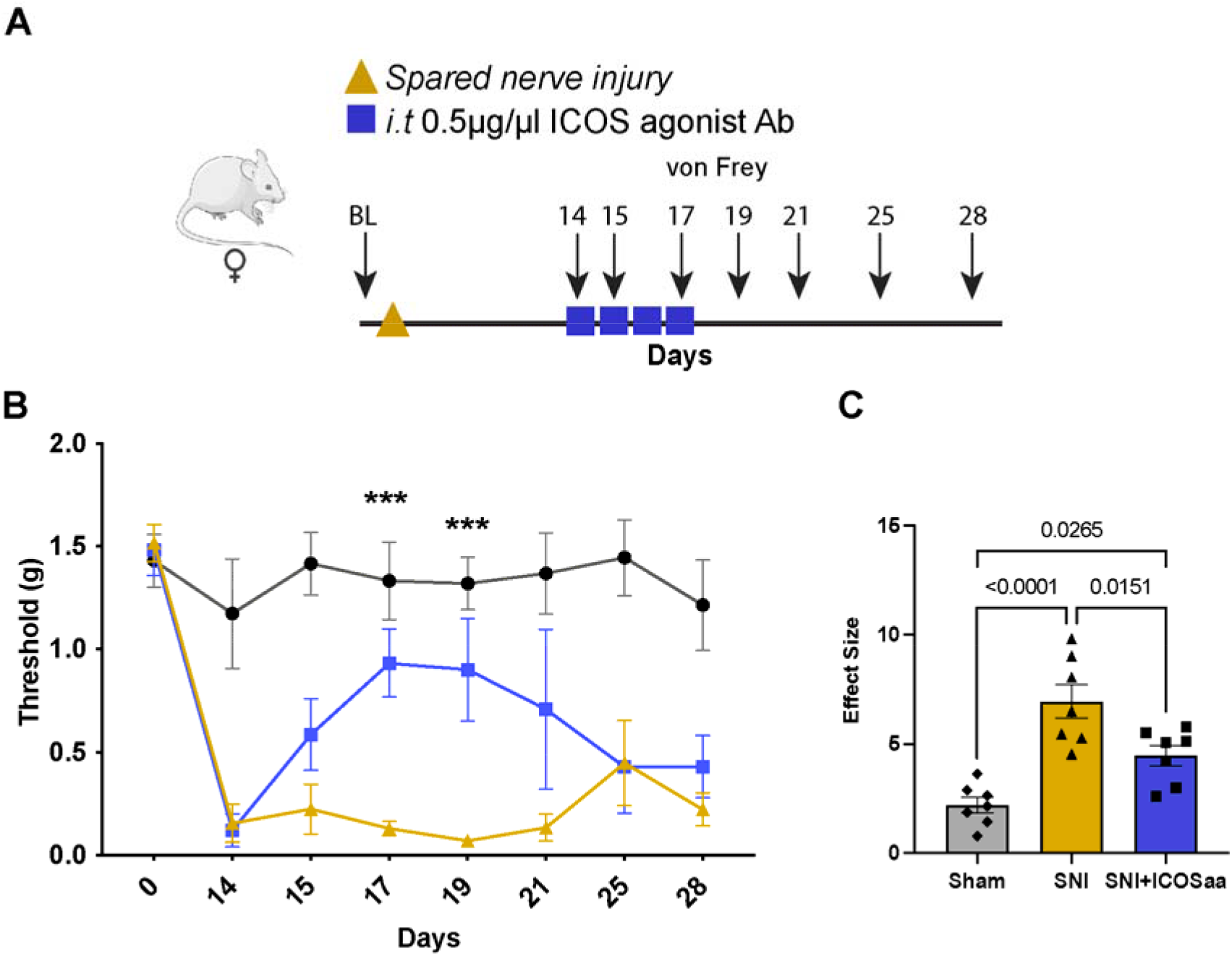
ICOSaa partially reverses mechanical allodynia in female SNI mice. A) Schematic representation of groups subjected to SNI on day one followed by intrathecal injection of ICOSaa on days 14-17. Arrows represent days with von Frey testing. B) Mechanical hypersentivity was partially reversed only on days 17 and 19 in the group administered with ICOSaa group (Two-way ANOVA, F=3.473, p-value<0.0001, post-hoc Bonferroni, ***SNI vs. SNI+ICOSaa at day 17, p-value=0.0006, ***SNI vs. SNI+ICOSaa at day 19, p-value=0.0003), N=7/group. C) Effect size was determined by calculating the cumulative difference between the value for each time point and the baseline value. The effect size was significant between SNI and SNI+ICOSaa treated mice (One-way ANOVA, F=18.0, p-value<0.0001, post-hoc Tukey, Sham vs. SNI+ICOSaa, p-value=0.0265, Sham vs. SNI, p=value<0.0001, SNI vs. SNI+ICOS, p-value=0.0151). ***p < 0.001.

### 3.5 Presence of T cells in human DRG

Our findings suggest that T cells in the DRG can be manipulated by ICOSaa treatment to increase IL-10 expression and alleviate neuropathic pain. To examine the translational potential of this approach, we investigated whether T cells are found in the human DRG using DRGs recovered from organ donors. We performed IHC using markers for CD4 and CD8a T cells. We observed that both CD4 and CD8a T cells were present in the DRG of organ donors, with many of these cells clustered around neurons (Fig 7). This is contrast with findings in naïve mouse DRG, where T cells are not present or are found only in low numbers [21]. This finding supports the translational potential of ICOSaa treatment for neuropathic pain.

**Figure 7:**
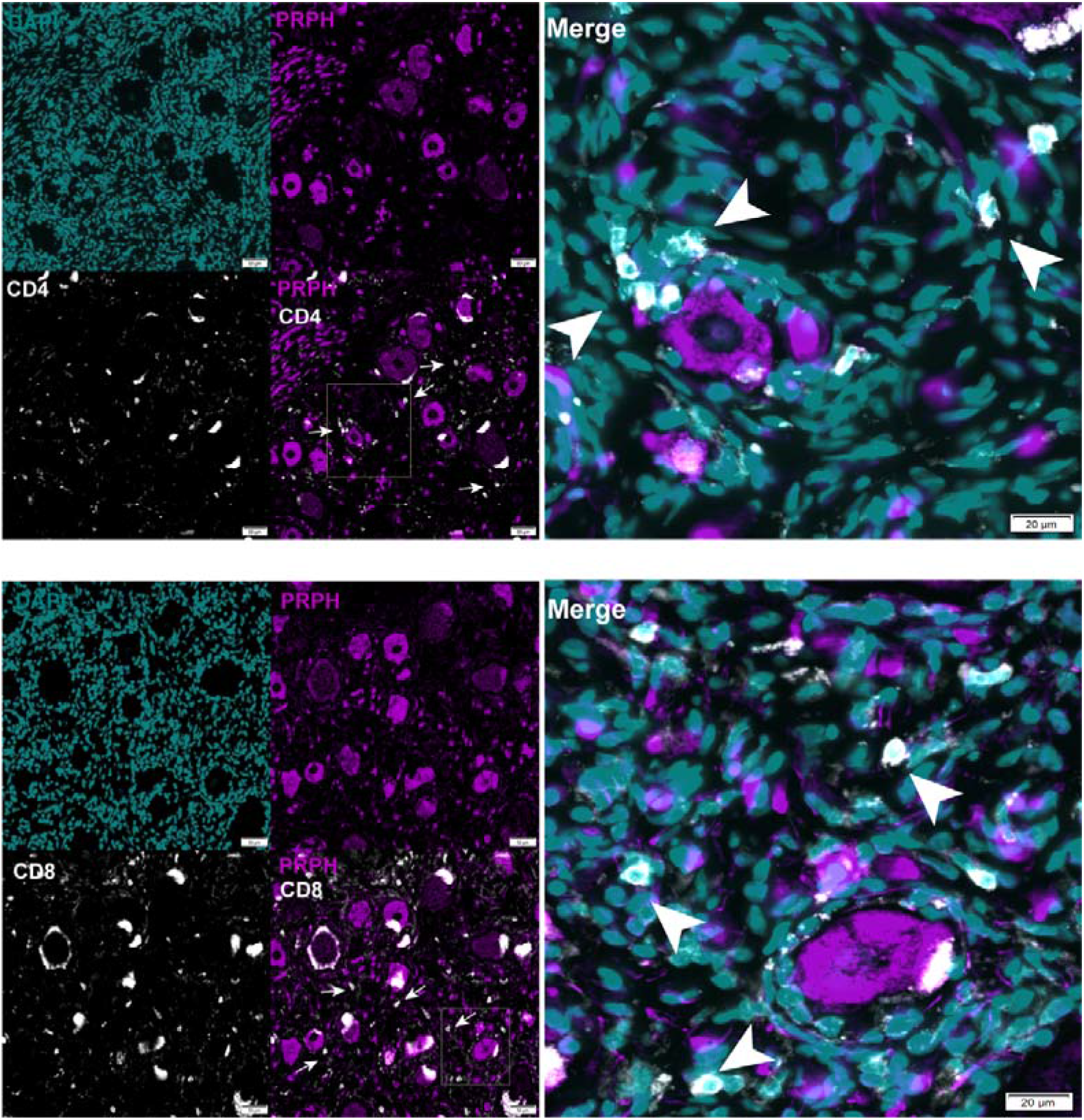
T cell expression in human DRG. A) representative image of CD4 (white) peripherin (purple), DAPI (blue) T cell expression in human dorsal root ganglion. B) representative image of CD8 (white) peripherin (purple), DAPI (cyan) T cell expression in human DRG. The white arrows point towards CD4 or CD8 T cell. Scale bar = 50 µm and 20 µm, respectively.

## Discussion

Our results support the hypothesis that targeting ICOS molecule facilitates the resolution of neuropathic pain in female mice. ICOS treatment enhanced the production of the anti-inflammatory cytokine IL-10 in the DRG of paclitaxel treated female mice, that subsequently, led to reversal of mechanical allodynia. Our findings are in line with literature that T cells can play a beneficial role in promoting neuropathic pain resolution [21; 24; 37]. Our study suggests a path for development of a completely new strategy for exogenously modulating T cells to facilitate pain resolution. Our work also shows that T cells infiltrate into the DRG after paclitaxel treatment. This is likely critical for pain resolution induced by ICOSaa as ICOS is thought to only be present on the surface of activated T cells [5; 27]. In a recent study, CD8 T cells were shown to play a key role in alleviating CIPN induced by cisplatin via an increase in the expression of IL-10R in the DRG [21; 23]. These observations are consistent with our results that administration of paclitaxel promotes T cell infiltration into the DRG and that ICOSaa treatment likely directs these cells towards a phenotype that promotes pain resolution.

ICOSaa treatment has been shown to mitigate disease severity in autoimmune disorders, lupus and cancers [2]. ICOS-ICOSL signaling has dual functions. On the one hand, it can suppress T regulatory cells and on the other it can cause T effector cells to express and secrete anti-inflammatory cytokines such as IL-4 and IL-10 [58]. Here we show that administration of ICOSaa increases anti-inflammatory cytokine IL-10 in the female mouse DRG. We suggest this administration activated the ICOS-ICOSL signaling cascade [10; 32; 48] to promote the release of anti-inflammatory, pro-pain resolution factors. ICOSaa treatment led to resolution of mechanical hypersensitivity in paclitaxel treated female mice and also trended towards the reversal of satellite cell gliosis in the DRG and reversed astrocyte gliosis in the dorsal horn of the spinal cord. This action on cellular measures of CIPN suggests the possibility of disease-modifying properties of ICOSaa treatment. Further studies will be needed to ascertain if ICOSaa treatment can also prevent the development of CIPN with paclitaxel treatment and/or if it is effective in other species. The observation of T cells in the DRGs of organ donors is promising from the perspective of clinical translation for these findings.

Cytokines are an important group of molecules that regulate the excitability of nociceptors and for this reason are intensively investigated for the treatment of pain where agonist and antagonist cytokine therapies are under development [17]. IL-10 has been shown to alleviate CIPN and rheumatoid arthritis [59]. *In vitro* studies show that ICOSaa activates the secretion of anti-inflammatory cytokines such as IL-4 and IL-10 [4]. Our results are in line with this literature as we have shown here that ICOSaa administration likely leads to secretion of the anti-inflammatory cytokine IL-10 in the DRG because expression of the cytokine was increased and effects of ICOSaa were blocked by IL-10 sequestering treatment. IL-10R is expressed on sensory neurons in the rodent DRG, and also on human DRG neurons [56], and its activation in rodent nociceptors reduces their excitability [22]. We are not aware of evidence that IL-10R is expressed by SGCs or astrocytes in the spinal cord, and single cell sequencing suggests that this receptor is exclusively found in neurons in the DRG [62]. Based on this, we speculate that the effect on astrogliosis and SGC gliosis is caused by the direct effects of IL-10 on DRG neurons. Reversing spontaneous activity in these neurons may allow for resolution of gliosis in the DRG and dorsal horn. Although IL-10 therapy can alleviate neuropathic pain, IL-10 has a short half-life under physiological conditions and the plasmid-based DNA therapy which showed promising results on canines with osteoarthritis is not yet FDA approved [53; 59]. Clinical trials with systemic administration of IL-10 did not downregulate inflammation due to the low concentration accumulated at the site of inflammation and higher dose IL-10 administration can be detrimental (Asadullah et al., 2003) Our hypothesized model of educating T cells to release IL-10 with ICOSaa may be able to overcome some of these limitations with a therapeutic approach that is nearing the clinic for other indications [54].

We observed a transient anti-nociceptive effect with the administration of ICOSaa in the SNI model of neuropathic pain. A possible explanation for this result could be that the mechanisms for the development and maintenance of pain in the SNI model is not entirely T cell dependent and that plasticity is developed with nerve injury. The role of T cells in trauma-induced neuropathic pain is still somewhat controversial. A previous study using the SNI model in Rag1^-/-^ mice (T cell deficient mice) reported less mechanical allodynia [9]. However, another study using the same model showed that T cells likely play a more important role in promoting neuropathic pain in female mice than in male mice [55]. Since this time, it has become clear that the type of T cell is critical for understanding the impact of these cells on neuropathic pain and neuropathic pain resolution [52]. Another potential reason for this difference in the duration of effect is that the SNI model creates a situation where nociceptor spontaneous activity is continuously driven by the ligation injury to the injured nerves. In the CIPN model the chemotherapy treatment is ceased prior to the ICOSaa treatment and there is no ongoing injury to drive the reemergence of neuropathic pain in this model following cessation of ICOSaa treatment.

A key finding of our study is the presence of CD8+ T cells in human DRG collected from healthy donors. This strengthens the case for the translational potential of ICOSaa. In naïve mice there are few T cells present in the DRG but the efficacy of ICOSaa treatment is presumably facilitated by the paclitaxel challenge promoting the infiltration of T cells into the DRG [21]. Our work demonstrates that both CD4+ and CD8+ T cells are present in human, healthy DRGs and are in the vicinity of neurons. To date, multiple Phase I and II clinical trials are being conducted for ICOS agonists and ICOS antagonists due to the high expression of ICOS in the tumor microenvironment and duality of function for the treatment of advance solid tumors [14; 25]. We think that our findings support the possibility that ICOS agonists could be used for pain treatment.

There are some limitations to the work described here. The first is that we cannot completely rule out a beneficial effect of ICOSaa treatment on male mice. In our study, paclitaxel treated male mice demonstrated a far more rapid resolution of mechanical hypersensitivity than female mice with vehicle treatment so it is difficult to ascertain whether there is a treatment effect or not over the time course of the experiment in males. The faster resolution of CIPN signs in these male mice creates a problem of statistical power as the expected effect size was smaller in this cohort of mice. Another weakness is that we have not tested the effect of ICOSaa treatment in mice depleted of T cells or looked at its effect on meningeal T regulatory cells.

Experiments using T cell depleted mice or examining meningeal immune cell populations could further delineate the mechanism of action of ICOSaa in neuropathic pain. Finally, our study was only done in mice. Replicating this work in another species and using human DRGs to better understand the clinical translational potential of ICOSaa treatment will be important to fully understand the translational potential of this novel approach.

## Acknowledgements

The authors thank the organ donors and their families for their enduring gift. We thank employees of the Southwest Transplant Alliance and members of the Price Lab for coordinating and assisting in DRGs recoveries, respectively.

## References Cited

[1] Agalave NM, Mody PH, Szabo-Pardi TA, Jeong HS, Burton MD. Neuroimmune Consequences of eIF4E Phosphorylation on Chemotherapy-Induced Peripheral Neuropathy. Front Immunol 2021;12:642420.

[2] Amatore F, Gorvel L, Olive D. Inducible Co-Stimulator (ICOS) as a potential therapeutic target for anti-cancer therapy. Expert Opin Ther Targets 2018;22(4):343–351.

[3] Amatore F, Gorvel L, Olive D. Role of Inducible Co-Stimulator (ICOS) in cancer immunotherapy. Expert Opin Biol Ther 2020;20(2):141–150.

[4] Arimura Y, Kato H, Dianzani U, Okamoto T, Kamekura S, Buonfiglio D, Miyoshi-Akiyama T, Uchiyama T, Yagi J. A co-stimulatory molecule on activated T cells, H4/ICOS, delivers specific signals in T(h) cells and regulates their responses. Int Immunol 2002;14(6):555–566.

[5] Austin PJ, Kim CF, Perera CJ, Moalem-Taylor G. Regulatory T cells attenuate neuropathic pain following peripheral nerve injury and experimental autoimmune neuritis. Pain 2012;153(9):1916–1931.

[6] Boyette-Davis JA, Cata JP, Driver LC, Novy DM, Bruel BM, Mooring DL, Wendelschafer-Crabb G, Kennedy WR, Dougherty PM. Persistent chemoneuropathy in patients receiving the plant alkaloids paclitaxel and vincristine. Cancer Chemother Pharmacol 2013;71(3):619–626.

[7] Brandolini L, d’Angelo M, Antonosante A, Allegretti M, Cimini A. Chemokine Signaling in Chemotherapy-Induced Neuropathic Pain. Int J Mol Sci 2019;20(12).

[8] Chaplan SR, Bach FW, Pogrel JW, Chung JM, Yaksh TL. Quantitative assessment of tactile allodynia in the rat paw. J Neurosci Methods 1994;53(1):55–63.

[9] Costigan M, Moss A, Latremoliere A, Johnston C, Verma-Gandhu M, Herbert TA, Barrett L, Brenner GJ, Vardeh D, Woolf CJ, Fitzgerald M. T-cell infiltration and signaling in the adult dorsal spinal cord is a major contributor to neuropathic pain-like hypersensitivity. J Neurosci 2009;29(46):14415–14422.

[10] Da Mesquita S, Papadopoulos Z, Dykstra T, Brase L, Farias FG, Wall M, Jiang H, Kodira CD, de Lima KA, Herz J, Louveau A, Goldman DH, Salvador AF, Onengut-Gumuscu S, Farber E, Dabhi N, Kennedy T, Milam MG, Baker W, Smirnov I, Rich SS, Dominantly Inherited Alzheimer N, Benitez BA, Karch CM, Perrin RJ, Farlow M, Chhatwal JP, Holtzman DM, Cruchaga C, Harari O, Kipnis J. Meningeal lymphatics affect microglia responses and anti-Abeta immunotherapy. Nature 2021;593(7858):255–260.

[11] Decosterd I, Woolf CJ. Spared nerve injury: an animal model of persistent peripheral neuropathic pain. Pain 2000;87(2):149–158.

[12] Duggett NA, Griffiths LA, McKenna OE, de Santis V, Yongsanguanchai N, Mokori EB, Flatters SJ. Oxidative stress in the development, maintenance and resolution of paclitaxel-induced painful neuropathy. Neuroscience 2016;333:13–26.

[13] Flatters SJ, Bennett GJ. Studies of peripheral sensory nerves in paclitaxel-induced painful peripheral neuropathy: evidence for mitochondrial dysfunction. Pain 2006;122(3):245–257.

[14] Garber K. Immune agonist antibodies face critical test. Nature reviews Drug discovery 2020;19(1):3–5.

[15] Gornstein E, Schwarz TL. The paradox of paclitaxel neurotoxicity: Mechanisms and unanswered questions. Neuropharmacology 2014;76 Pt A:175–183.

[16] Hylden JL, Wilcox GL. Intrathecal morphine in mice: a new technique. European journal of pharmacology 1980;67(2-3):313–316.

[17] Inyang KE, Folger JK, Laumet G. Can FDA-Approved Immunomodulatory Drugs be Repurposed/Repositioned to Alleviate Chronic Pain? J Neuroimmune Pharmacol 2021;16(3):531–547.

[18] Iyer SS, Cheng G. Role of interleukin 10 transcriptional regulation in inflammation and autoimmune disease. Crit Rev Immunol 2012;32(1):23–63.

[19] Ji RR, Chamessian A, Zhang YQ. Pain regulation by non-neuronal cells and inflammation. Science 2016;354(6312):572–577.

[20] Krishnan AV, Goldstein D, Friedlander M, Kiernan MC. Oxaliplatin-induced neurotoxicity and the development of neuropathy. Muscle Nerve 2005;32(1):51–60.

[21] Krukowski K, Eijkelkamp N, Laumet G, Hack CE, Li Y, Dougherty PM, Heijnen CJ, Kavelaars A. CD8+ T Cells and Endogenous IL-10 Are Required for Resolution of Chemotherapy-Induced Neuropathic Pain. J Neurosci 2016;36(43):11074–11083.

[22] Laumet G, Bavencoffe A, Edralin JD, Huo XJ, Walters ET, Dantzer R, Heijnen CJ, Kavelaars A. Interleukin-10 resolves pain hypersensitivity induced by cisplatin by reversing sensory neuron hyperexcitability. Pain 2020;161(10):2344–2352.

[23] Laumet G, Edralin JD, Dantzer R, Heijnen CJ, Kavelaars A. Cisplatin educates CD8+ T cells to prevent and resolve chemotherapy-induced peripheral neuropathy in mice. Pain 2019;160(6):1459–1468.

[24] Laumet G, Ma J, Robison AJ, Kumari S, Heijnen CJ, Kavelaars A. T Cells as an Emerging Target for Chronic Pain Therapy. Front Mol Neurosci 2019;12:216.

[25] Le Tourneau C, Rischin D, Groenland S, Lim A, Martin-Liberal J, Moreno V, Trigo J, Mathew M, Cho D, Hansen A. 1O Inducible T cell co-stimulatory (ICOS) receptor agonist, GSK3359609 (GSK609) alone and combination with pembrolizumab: Preliminary results from INDUCE-1 expansion cohorts in head and neck squamous cell carcinoma (HNSCC). Annals of Oncology 2020;31:S1.

[26] Ledeboer A, Jekich BM, Sloane EM, Mahoney JH, Langer SJ, Milligan ED, Martin D, Maier SF, Johnson KW, Leinwand LA, Chavez RA, Watkins LR. Intrathecal interleukin-10 gene therapy attenuates paclitaxel-induced mechanical allodynia and proinflammatory cytokine expression in dorsal root ganglia in rats. Brain Behav Immun 2007;21(5):686–698.

[27] Lees JG, Duffy SS, Perera CJ, Moalem-Taylor G. Depletion of Foxp3+ regulatory T cells increases severity of mechanical allodynia and significantly alters systemic cytokine levels following peripheral nerve injury. Cytokine 2015;71(2):207–214.

[28] Lees JG, Makker PG, Tonkin RS, Abdulla M, Park SB, Goldstein D, Moalem-Taylor G. Immune-mediated processes implicated in chemotherapy-induced peripheral neuropathy. Eur J Cancer 2017;73:22–29.

[29] Leo M, Schmitt L-I, Kutritz A, Kleinschnitz C, Hagenacker T. Cisplatin-induced activation and functional modulation of satellite glial cells lead to cytokine-mediated modulation of sensory neuron excitability. Experimental Neurology 2021;341:113695.

[30] Liu XJ, Zhang Y, Liu T, Xu ZZ, Park CK, Berta T, Jiang D, Ji RR. Nociceptive neurons regulate innate and adaptive immunity and neuropathic pain through MyD88 adapter. Cell Res 2014;24(11):1374–1377.

[31] Lo□hning M, Hutloff A, Kallinich T, Mages HW, Bonhagen K, Radbruch A, Hamelmann E, Kroczek RA. Expression of ICOS in vivo defines CD4+ effector T cells with high inflammatory potential and a strong bias for secretion of interleukin 10. The Journal of experimental medicine 2003;197(2):181–193.

[32] Louveau A, Herz J, Alme MN, Salvador AF, Dong MQ, Viar KE, Herod SG, Knopp J, Setliff JC, Lupi AL, Da Mesquita S, Frost EL, Gaultier A, Harris TH, Cao R, Hu S, Lukens JR, Smirnov I, Overall CC, Oliver G, Kipnis J. CNS lymphatic drainage and neuroinflammation are regulated by meningeal lymphatic vasculature. Nat Neurosci 2018;21(10):1380–1391.

[33] Maeda S, Fujimoto M, Matsushita T, Hamaguchi Y, Takehara K, Hasegawa M. Inducible costimulator (ICOS) and ICOS ligand signaling has pivotal roles in skin wound healing via cytokine production. The American journal of pathology 2011;179(5):2360–2369.

[34] Mahajan S, Cervera A, MacLeod M, Fillatreau S, Perona-Wright G, Meek S, Smith A, MacDonald A, Gray D. The role of ICOS in the development of CD4 T cell help and the reactivation of memory T cells. European journal of immunology 2007;37(7):1796–1808.

[35] Makker PG, Duffy SS, Lees JG, Perera CJ, Tonkin RS, Butovsky O, Park SB, Goldstein D, Moalem-Taylor G. Characterisation of immune and neuroinflammatory changes associated with chemotherapy-induced peripheral neuropathy. PloS one 2017;12(1):e0170814.

[36] Malacrida A, Meregalli C, Rodriguez-Menendez V, Nicolini G. Chemotherapy-induced peripheral neuropathy and changes in cytoskeleton. International journal of molecular sciences 2019;20(9):2287.

[37] Mao L, Li P, Zhu W, Cai W, Liu Z, Wang Y, Luo W, Stetler RA, Leak RK, Yu W, Gao Y, Chen J, Chen G, Hu X. Regulatory T cells ameliorate tissue plasminogen activator-induced brain haemorrhage after stroke. Brain 2017;140(7):1914–1931.

[38] McWhinney SR, Goldberg RM, McLeod HL. Platinum neurotoxicity pharmacogenetics. Mol Cancer Ther 2009;8(1):10–16.

[39] Megat S, Ray PR, Moy JK, Lou TF, Barragan-Iglesias P, Li Y, Pradhan G, Wanghzou A, Ahmad A, Burton MD, North RY, Dougherty PM, Khoutorsky A, Sonenberg N, Webster KR, Dussor G, Campbell ZT, Price TJ. Nociceptor Translational Profiling Reveals the Ragulator-Rag GTPase Complex as a Critical Generator of Neuropathic Pain. J Neurosci 2019;39(3):393–411.

[40] Milligan ED, Sloane EM, Langer SJ, Hughes TS, Jekich BM, Frank MG, Mahoney JH, Levkoff LH, Maier SF, Cruz PE. Repeated intrathecal injections of plasmid DNA encoding interleukin-10 produce prolonged reversal of neuropathic pain. Pain 2006;126(1-3):294–308.

[41] Ng TH, Britton GJ, Hill EV, Verhagen J, Burton BR, Wraith DC. Regulation of adaptive immunity; the role of interleukin-10. Front Immunol 2013;4:129.

[42] Park SB, Goldstein D, Krishnan AV, Lin CS, Friedlander ML, Cassidy J, Koltzenburg M, Kiernan MC. Chemotherapy-induced peripheral neurotoxicity: a critical analysis. CA Cancer J Clin 2013;63(6):419–437.

[43] Pekny M, Pekna M. Astrocyte reactivity and reactive astrogliosis: costs and benefits. Physiological reviews 2014;94(4):1077–1098.

[44] Peters CM, Jimenez-Andrade JM, Jonas BM, Sevcik MA, Koewler NJ, Ghilardi JR, Wong GY, Mantyh PW. Intravenous paclitaxel administration in the rat induces a peripheral sensory neuropathy characterized by macrophage infiltration and injury to sensory neurons and their supporting cells. Exp Neurol 2007;203(1):42–54.

[45] Prado J, Westerink RH, Popov-Celeketic J, Steen-Louws C, Pandit A, Versteeg S, van de Worp W, Kanters DH, Reedquist KA, Koenderman L. Cytokine receptor clustering in sensory neurons with an engineered cytokine fusion protein triggers unique pain resolution pathways. Proceedings of the National Academy of Sciences 2021;118(11).

[46] Robinson CR, Zhang H, Dougherty PM. Astrocytes, but not microglia, are activated in oxaliplatin and bortezomib-induced peripheral neuropathy in the rat. Neuroscience 2014;274:308–317.

[47] Sahenk Z, Barohn R, New P, Mendell JR. Taxol neuropathy. Electrodiagnostic and sural nerve biopsy findings. Arch Neurol 1994;51(7):726–729.

[48] Salvador AF, de Lima KA, Kipnis J. Neuromodulation by the immune system: a focus on cytokines. Nat Rev Immunol 2021;21(8):526–541.

[49] Seretny M, Currie GL, Sena ES, Ramnarine S, Grant R, MacLeod MR, Colvin LA, Fallon M. Incidence, prevalence, and predictors of chemotherapy-induced peripheral neuropathy: A systematic review and meta-analysis. Pain 2014;155(12):2461–2470.

[50] Shen K-F, Zhu H-Q, Wei X-H, Wang J, Li Y-Y, Pang R-P, Liu X-G. Interleukin-10 down-regulates voltage gated sodium channels in rat dorsal root ganglion neurons. Experimental neurology 2013;247:466–475.

[51] Singh AK, Mahalingam R, Squillace S, Jacobson KA, Tosh DK, Dharmaraj S, Farr SA, Kavelaars A, Salvemini D, Heijnen CJ. Targeting the A3 adenosine receptor to prevent and reverse chemotherapy-induced neurotoxicities in mice. Acta Neuropathol Commun 2022;10(1):11.

[52] Singh SK, Krukowski K, Laumet GO, Weis D, Alexander JF, Heijnen CJ, Kavelaars A. CD8+ T cell–derived IL-13 increases macrophage IL-10 to resolve neuropathic pain. JCI insight 2022;7(5).

[53] Soderquist RG, Milligan ED, Harrison JA, Chavez RA, Johnson KW, Watkins LR, Mahoney MJ. PEGylation of interleukin-10 for the mitigation of enhanced pain states. J Biomed Mater Res A 2010;93(3):1169–1179.

[54] Solinas C, Gu-Trantien C, Willard-Gallo K. The rationale behind targeting the ICOS-ICOS ligand costimulatory pathway in cancer immunotherapy. ESMO Open 2020;5(1).

[55] Sorge RE, Mapplebeck JC, Rosen S, Beggs S, Taves S, Alexander JK, Martin LJ, Austin JS, Sotocinal SG, Chen D, Yang M, Shi XQ, Huang H, Pillon NJ, Bilan PJ, Tu Y, Klip A, Ji RR, Zhang J, Salter MW, Mogil JS. Different immune cells mediate mechanical pain hypersensitivity in male and female mice. Nat Neurosci 2015;18(8):1081–1083.

[56] Tavares-Ferreira D, Shiers S, Ray PR, Wangzhou A, Jeevakumar V, Sankaranarayanan I, Cervantes AM, Reese JC, Chamessian A, Copits BA, Dougherty PM, Gereau RWt, Burton MD, Dussor G, Price TJ. Spatial transcriptomics of dorsal root ganglia identifies molecular signatures of human nociceptors. Sci Transl Med 2022;14(632):eabj8186.

[57] Tonini G, Santini D, Vincenzi B, Borzomati D, Dicuonzo G, La Cesa A, Onori N, Coppola R. Oxaliplatin may induce cytokine-release syndrome in colorectal cancer patients. Journal of biological regulators and homeostatic agents 2002;16(2):105–109.

[58] Watanabe M, Takagi Y, Kotani M, Hara Y, Inamine A, Hayashi K, Ogawa S, Takeda K, Tanabe K, Abe R. Down-regulation of ICOS ligand by interaction with ICOS functions as a regulatory mechanism for immune responses. J Immunol 2008;180(8):5222–5234.

[59] Watkins LR, Chavez RA, Landry R, Fry M, Green-Fulgham SM, Coulson JD, Collins SD, Glover DK, Rieger J, Forsayeth JR. Targeted interleukin-10 plasmid DNA therapy in the treatment of osteoarthritis: Toxicology and pain efficacy assessments. Brain Behav Immun 2020;90:155–166.

[60] Wikenheiser DJ, Stumhofer JS. ICOS Co-Stimulation: Friend or Foe? Front Immunol 2016;7:304.

[61] Yanik BM, Dauch JR, Cheng HT. Interleukin-10 reduces neurogenic inflammation and pain behavior in a mouse model of type 2 diabetes. Journal of pain research 2020;13:3499.

[62] Zeisel A, Hochgerner H, Lonnerberg P, Johnsson A, Memic F, van der Zwan J, Haring M, Braun E, Borm LE, La Manno G, Codeluppi S, Furlan A, Lee K, Skene N, Harris KD, Hjerling-Leffler J, Arenas E, Ernfors P, Marklund U, Linnarsson S. Molecular Architecture of the Mouse Nervous System. Cell 2018;174(4):999-1014.e1022.

[63] Zhang H, Li Y, de Carvalho-Barbosa M, Kavelaars A, Heijnen CJ, Albrecht PJ, Dougherty PM. Dorsal Root Ganglion Infiltration by Macrophages Contributes to Paclitaxel Chemotherapy-Induced Peripheral Neuropathy. J Pain 2016;17(7):775–786.

